# Mechanical management of weeds drops nymphal density of *Xylella fastidiosa* vectors

**DOI:** 10.1101/2022.10.18.512680

**Authors:** Júlia López-Mercadal, Pau Mercadal-Frontera, Miguel Ángel Miranda

## Abstract

*Xylella fastidiosa* Wells (1987) (Proteobacteria:Xanthomonadaceae) is a xylem pathogen bacterium transmitted by xylem feeder insects that causes several important plant diseases such as Pierce’s disease in grapes or leaf scorch in almond and olives trees. The bacterium was detected in the Balearic Islands in October 2016, including three subspecies: *fastidiosa*, *multiplex* and *pauca*. The major potential vectors described in the Balearics are *Philaenus spumarius* L. and *Neophilaenus campestris* Fallen (1805). In order to interfere the life cycle of vectors, we tested the effect of mechanical control of the plant cover on the most vulnerable phases, such as nymphs and/or newly emerged adults. For this, we selected four organic orchards in Mallorca, three olive and one vineyard plots. Owners of each selected plot conducted mechanical control according to their common procedures and their own machinery, which in general included cut and tillage of the plant cover during March-April. Nymph abundance per surface (30 sampling points/treatment/orchard x 0,25 m2) was measured in each plot in a weekly basis before and after mechanical control. Our results indicated that either tillage and mowing decreased nymphal density of *X. fastidiosa* vectors in both types of crops. These results contribute to the integrated pest management of vectors by conducting feasible farm-based management of the regular plant cover.

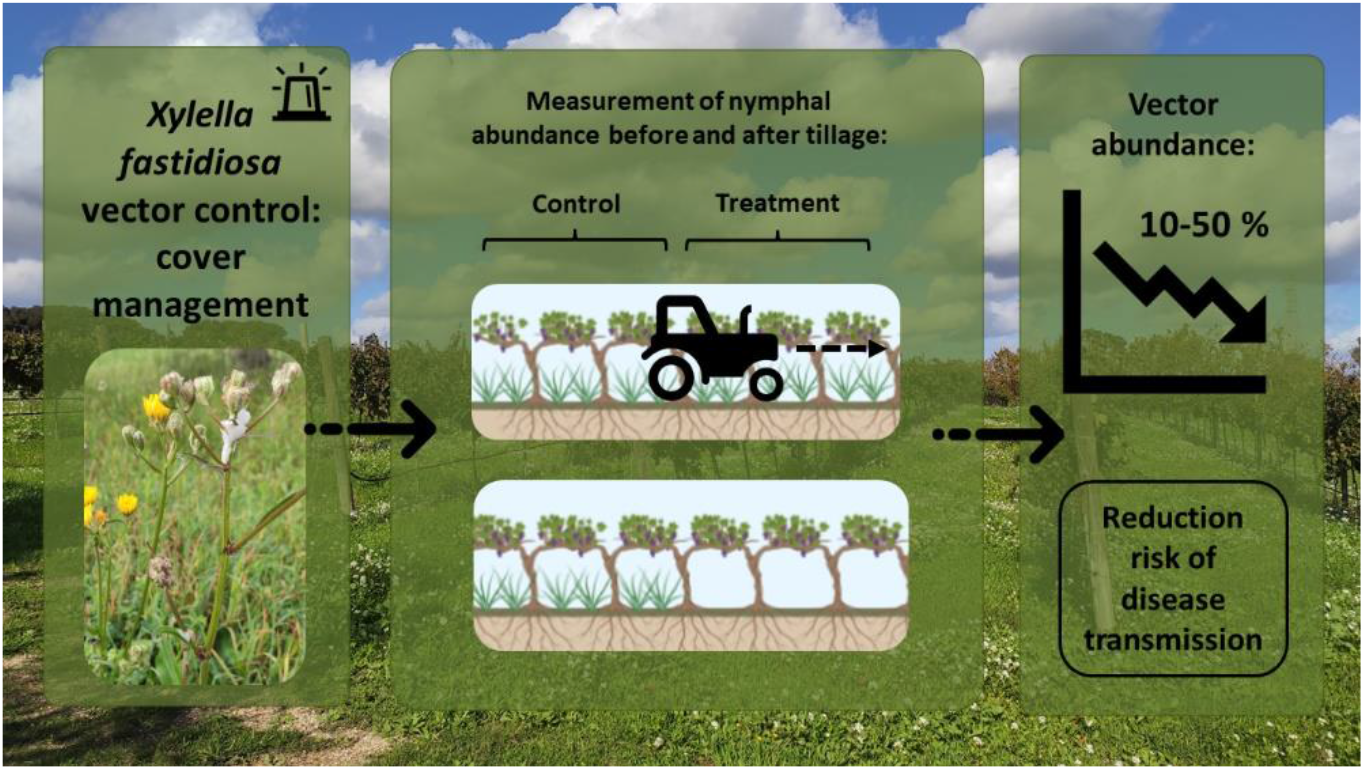

## 1. Introduction

Land-management strategies with environmental, social and economic benefits also include to maintain local biodiversity and associated ecosystem services such as pollination and pest control [1]. An effect of cover cropping is the increasing of biodiversity, to reduce number of specialized parasites and to increase ecological stability [2–5]. Intensive tillage has been shown to decrease plant and animal species diversity for some taxa [6–8], while the use of cover crops in vineyard inter-rows has showed to have positive effects on pest control [9–10], such as increasing food web complexity and intraguild predation [11]. On the contrary, certain plant species may also increase potential pest species by acting as a host plant [12], by providing resources or shelter [13]. There are different ways to manage weeds by the farmer such as flaming, mowing or tillage as an alternative to herbicides that are producing resistance in weed communities [14–15].

Cover plants are essential for the development of groups of insects such as Cicadomorpha (Hemiptera) [16–17], which are vectors of important pathogens such as *Xylella fastidiosa* Wells (1987) (Proteobacteria:Xanthomonadaceae) [18]. This bacterium is a pathogen of plants limited to the xylem and capable of infecting more than 600 plant species [19–21]. This species has great number of genotypic and phenotypic diversity, that allows the bacterium to have a wide host range [19–20,22–23]. Transmission of *X. fastidiosa* is conducted by the xylem feeding activity of Cicadomorpha adults, depending on its presence, abundance and behavior [24–29]. The bacterium is restricted to their alimentary canal, where they adhere to, multiply and persist in the precibarium and cibarium foregut parts of the insect [20,27].

Within Cicadomorpha, Aphrophoridae are the major vectors of *X. fastidiosa* in Europe [30]. *Philaenus spumarius* L., *P. italosignus* Drosopoulos and Remane (2000) and *Neophilaenus campestris* Fallen (1805) are considered proven vectors in Europe [30]. In the Balearic Islands (Spain), theyusually overwinter as egg form until March when nymphs start to emerge in the cover vegetation. Adults start to appear in end-April and remain in the cover until it dries in summer to migrate to tree canopies and bordering woody shrubs [31–32]. Then, adults return to cover in autumn for mating, completing their univoltine life cycle [32]. The bacterium is associated with important diseases in a wide range of plants, being an important emerging pathogen [26,33]. Each subspecies and genetic type (ST) have different host range causing diseases such as the Pierce’s disease in grapevine (*Vitis vinifera* L.), citrus variegated chlorosis, leaf scorch (almond, elm, oak, oleander, American sycamore, mulberry and maple), alfalfa dwarf, olive quick decline, plum leaf scald and peach phony rickettsia [28, 30, 34–36]. Nevertheless, many species of plants may remain symptomless [20,32]. In Europe, there are hosts with a high economic value such as *Olea europaea* L., *Prunus dulcis* (Mill.) D. A. Webb, *V. vinifera*, *P. avium* L., *P. domestica* L., *P. salicina* Lindl. or *Citrus* spp. L. [37], being olive and vineyard crops the largest cultivation in the Mediterranean basin [5].

*Xylella fastidiosa* vectors in Europe are not major pest and they usually cause little damage to plants [27]. Nevertheless, if *X. fastidiosa* was fully spread, it would cause an annual production loss of 5.5 billion euros that affects the 70 % of older olive trees (over 30 years old) and the 35 % of younger olive production; 13 % of almond, 11 % of citrus and 1-2 % of grapevine [38]. In Italy, it was estimated that olive producers have already lost between 0.2 and 0.6 billion euros in investments, and it could increase until 1.9 to 5.5 billion of euros over the next 50 years [39–40].

Since *X. fastidiosa* was first detected in Europe in 2013 [41], a huge effort was made to avoid the spread of the disease. Chemical curative control against the bacterium is still under study as zinc-copper-citric acid biocomplex has been shown to decrease bacterium population in olive trees [42–43]. Otherwise prevention by use of resistant varieties, hygienic and cultural measures (i.e., cover plant management), biological (i.e., parasitoids or spiders) and chemical (neonicotinoids and pyrethroids) vector control are the pathways to achieve it [35, 37]. Combining multiple of these control strategies is considered as the best management strategy [27]. But considering the negative effects of pesticides in the agrosystems, it is of major importance to develop more sustainable and eco-friendly control methods. The aim of this study is to assess the efficacy of mechanical control methods against *X. fastidiosa* nymphal vectors in olive and vine organic orchards in Majorca, considered to be of high economic value in the area.

## 2. Materials and Methods

### Study site

Three olive orchards were selected from Majorca (Balearic Islands), all of them under official organic farming management (Fig. 1). The climate in the Balearics in the Mediterranean type, characterized by dry and hot summers and wet mild winters. The annual mean temperature is 21.8 °C and the annual mean precipitation 456 mm [44].

**Figure 1.**
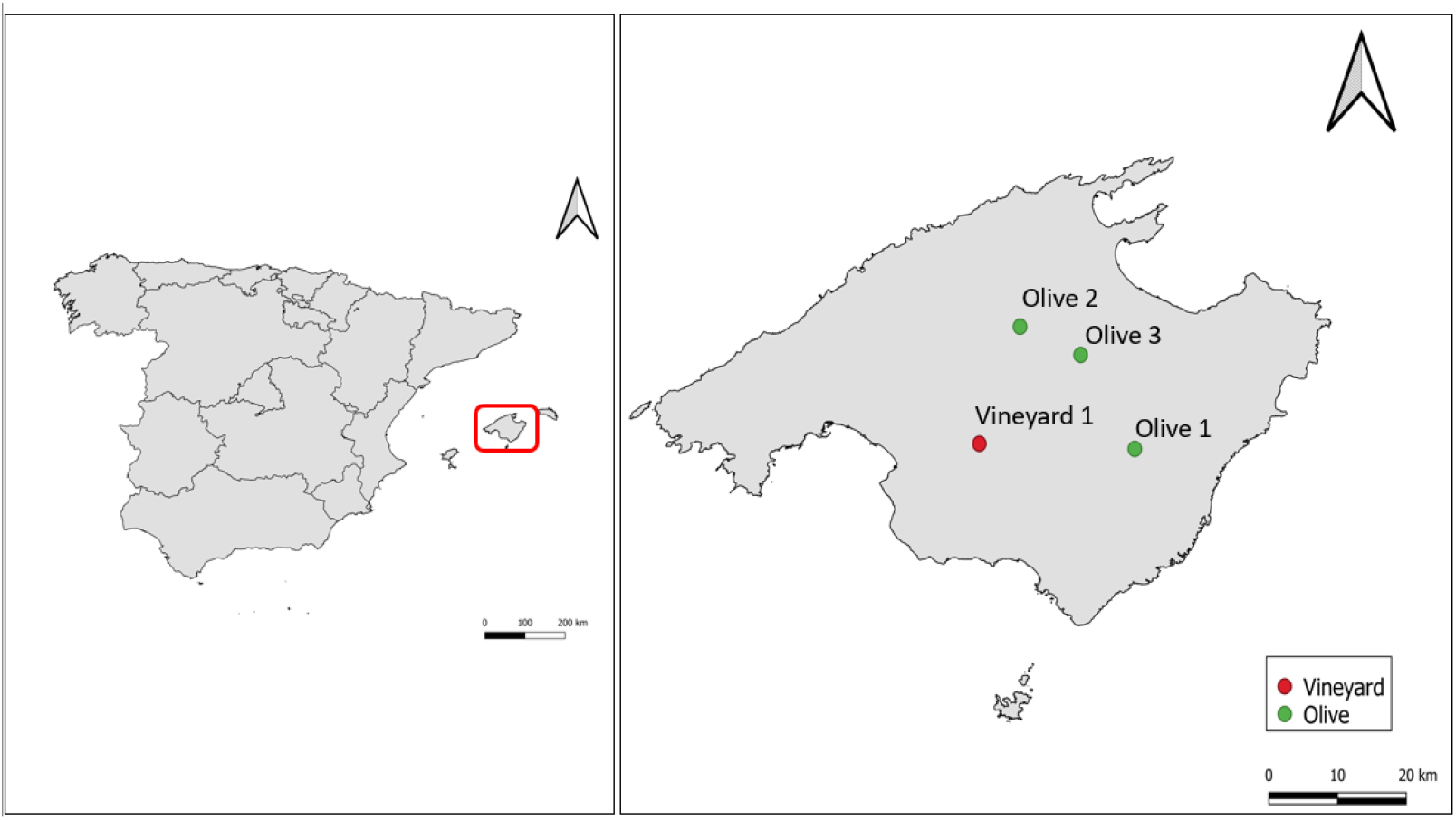
Location of the plots used for the trials. Olive plots are highlighted with red triangles and the vineyard plot with a red square. Olive 1: 39°31’39.31” N; 3° 8’58.71” E, 1.93 Ha; Olive 2: 39°42’34.80” N, 2°57’26.08” E, 2.66 Ha; Olive 3: 39°40’3.49” N, 3° 3’31.52” E, 0.26 Ha; Vineyard 1: 39°32’7.22” N; 2°53’20.42” E, 3.68 Ha. Mechanical control of nymphs in the cover vegetation.

The same methodology was used for olive and vineyard crops in 2020 and 2021. Nymphs of the vectors of *X. fastidiosa* were surveyed by visual counts from end-March in each plot. When nymphs were present, total density was determined and rows in the crop were marked as control or treatment (Fig. 2, Table 1). Density of nymphs was determined by using a 0.25 m^2^ woody rectangle. In each control and treatment rows, 30 samples per week were collected using the rectangle (90 samples/week/plot). Samples were collected weekly before treatment and two weeks after treatment. In each of the control and treatment rows in all the three samplings conducted following nymphal sampling methodology described in López-Mercadal et al., [32]. The position of each rectangle was marked with a rope to assess the density in the same place every time. We got a short time to conduct the experiments as nymphal period lasts from March to end May in Majorca and considering stages development among time [32], we decided to conduct three samplings. After first day of nymph sampling, cover vegetation was cut by farmers using regular equipment in the treatment rows. Hereafter, nymphal density was checked after 2 and 3 weeks from the tillage date in both treatment and control rows. Weed control in olive orchards were by mowing and in vineyard orchards by tillage. In the treated rows, ground vegetation never reached zero vegetation due to was not possible to till and cut near the trunks to not damage the olives and vine plants.

**Figure 2.**
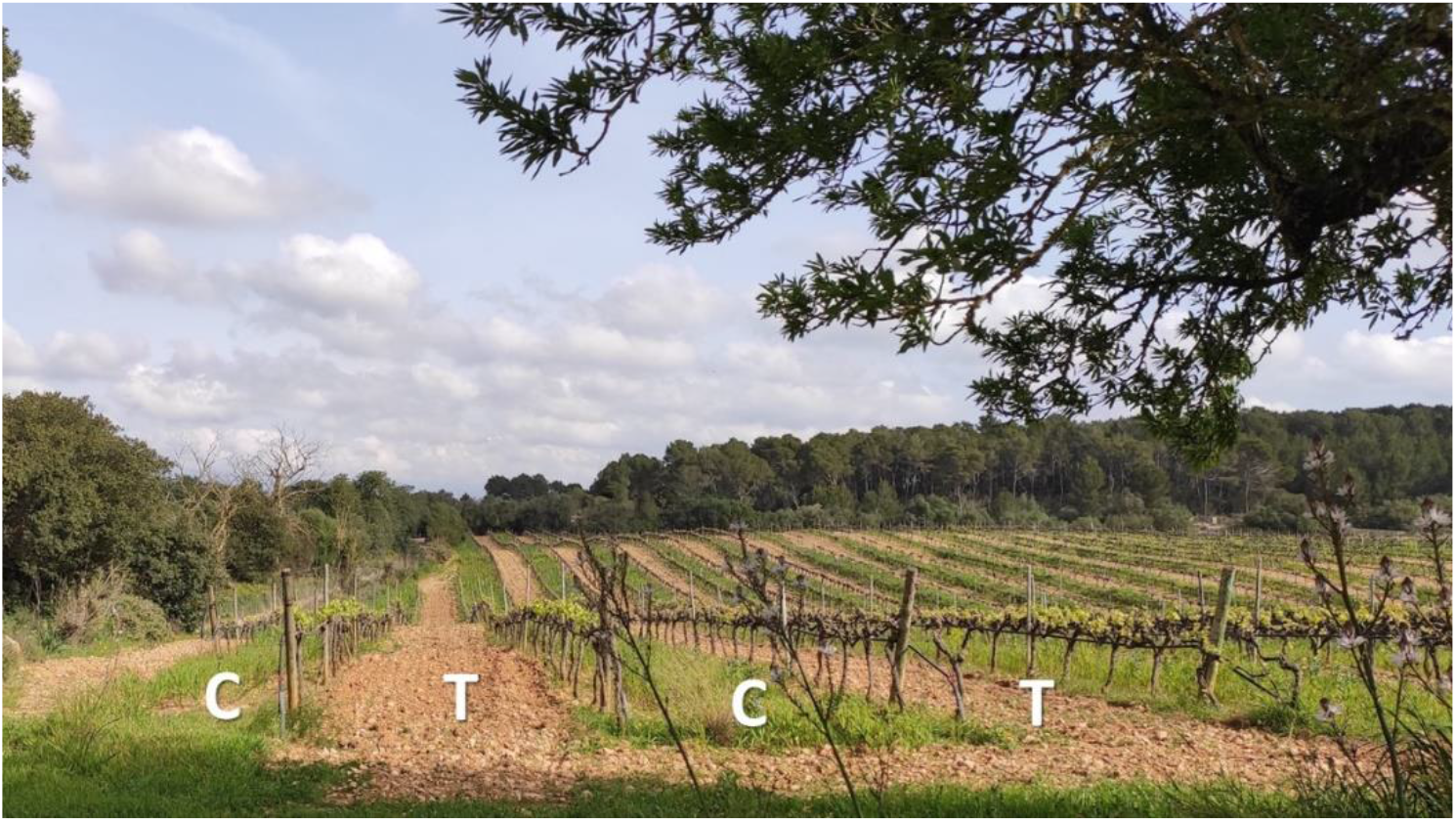
Example of experimental design conducted in a vineyard plot. Treatment row (T) where cover plants were mowed and control (C) rows where no tillage was conducted in a vineyard.

**Table 1.**
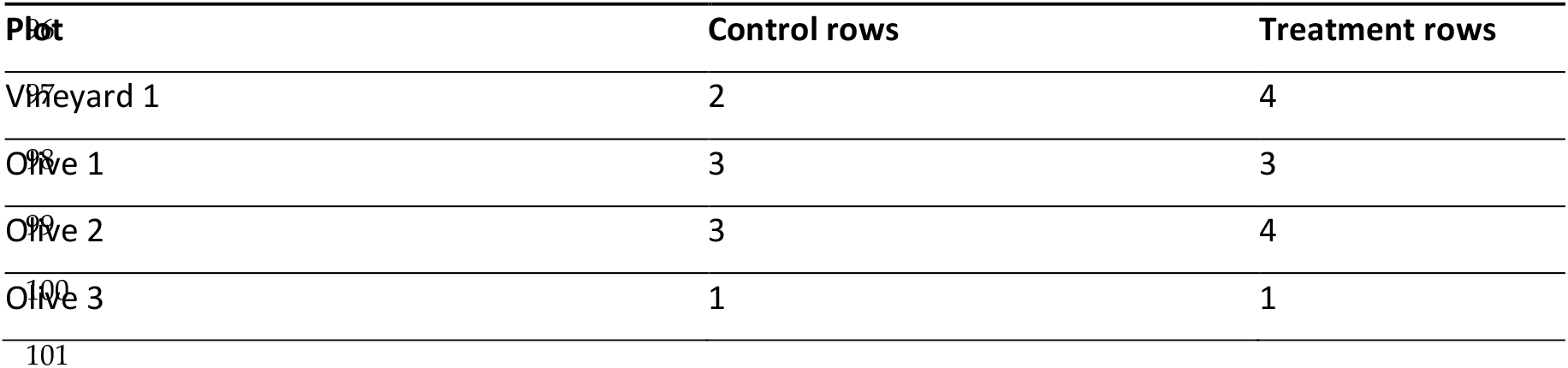
Number of rows tested in the trials in olive and vineyard.

| Plot | Control rows | Treatment rows |
| --- | --- | --- |
| Vineyard 1 | 2 | 4 |
| Olive 1 | 3 | 3 |
| Olive 2 | 3 | 4 |
| Olive 3 | 1 | 1 |

### Statistical analysis

Zero-inflated models were used to assess the influence of tillage or mowing on the cover herbaceous vegetation on nymphal density with the package pscl [45]. Week (1, 2 or 3), treatment (control or treatment) and time (pre-treatment or post-treatment) were included as fixed factors. Post-hoc analysis with Tukey adjustment were performed with emmeans [46] and multcomp packages [47]. We accepted as significant the p-values below 0.05. Statistical analyses were performed in R software 3.2.5 [48].

## 3. Results

### 2020 trials

The trial was conducted in one plot of olive and one of vineyard. In both plots there were nymphs of *P. spumarius* and *N. campestris*. In the case of the vine crop, initial density was the same for both zones with 0.2 nymphs/m^2^ (Estimate: 0.5839, Std. Error: 0.6865, P-value = 0.395) (Fig. 3 a). Due to zero density in the treatment zone in the first week post-treatment, data was not able to be analysed statistically. In the second week post-treatment we observed significantly differences among control and treatment (Estimate: −2.6135, Std. Error: 0.7119, P-value <0.05). In the case of olive crop (Fig. 3 b), initial density was the same for both zones with 0.35 nymphs/m^2^ (Estimate: −0.4170, Std. Error: 0.3844, P-value = 0.395). In the second week, we observed statistically differences among control and treatment (Estimate: 3.262, Std. Error: 1.302, P-value = <0.05). Finally, in the third week nymph density in the treatment rows (0.1 nymphs/m^2^) was lower than in the control rows (0.4 nymphs/m^2^), however, there was not significant difference between them (Estimate: −0.3860, Std. Error: 0.5156, P-value = 0.4541).

**Figure 3.**
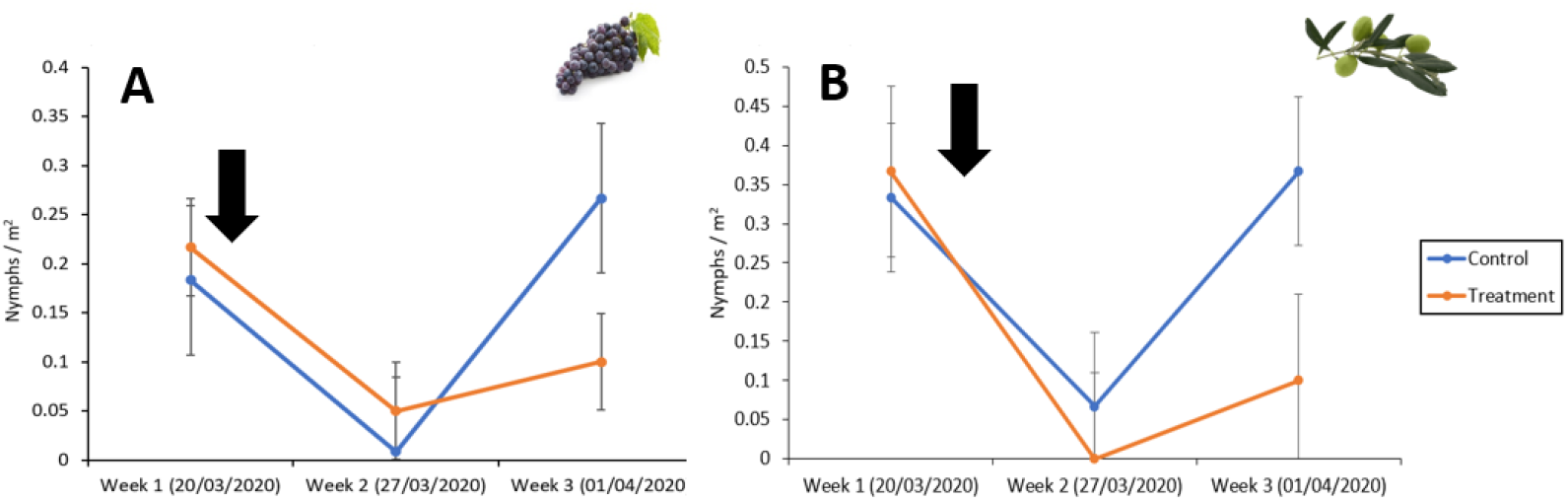
Nymph density per m^2^ ± SE (bars) in vineyard plot 1 (A) and in olive plot 1 (B) crops. Pre-treatment corresponds to week 1 and post-treatment to week 2 and 3. The black row indicates the moment tillage or mowing.

### 2021 trials

The trial was conducted in two organic olive orchards. In both plots there were nymphs of *P. spumarius* and *N. campestris*. Due to the difference of the dynamics of both plots, we decided to analyse them for separate. Initial density before tillage was statistically the same for treatment and control in the plot A (Estimate: 0.3622; Std. Error: 0.2587; P-value = 0.1615) (Fig. 4 a) as well as in the plot B (Estimate: 0.3622; Std. Error: 0.2587; P-value= 0.1615) (Figure 4 b).

**Figure 4.**
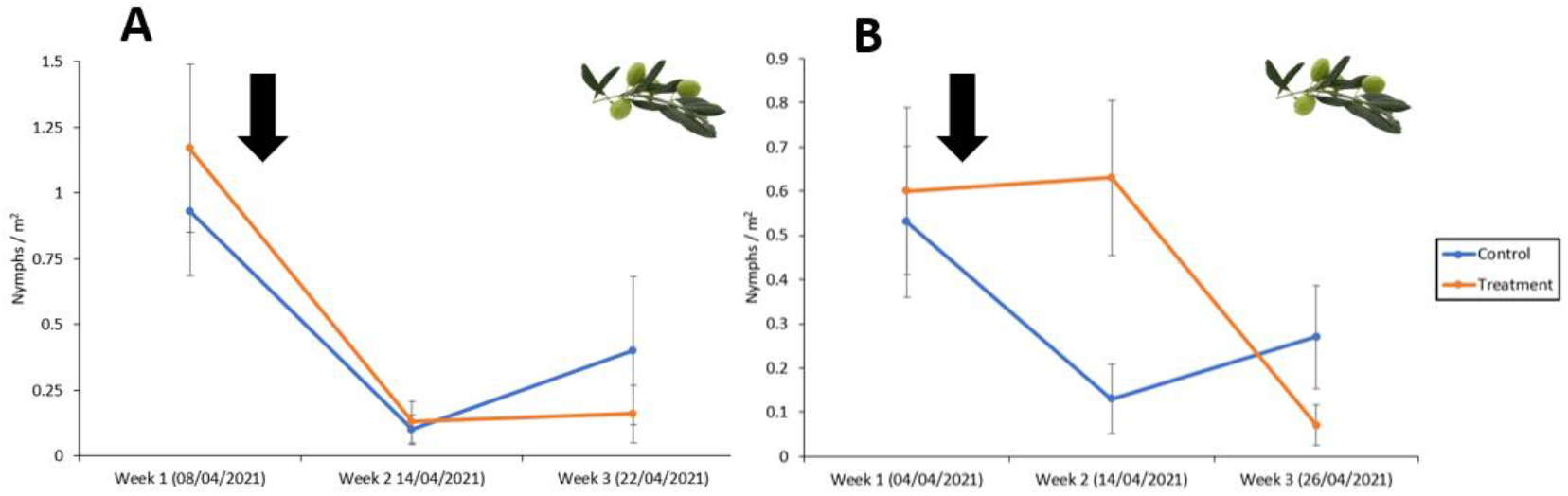
Nymph density per m^2^ ± SE (bars) in olive plot 2 and 3 (**A** and **B**, respectively). Pre-treatment corresponds to week 1 and post-treatment to week 2 and 3. The black row indicates the moment tillage or mowing.

In the plot A, after the tillage, there was no overall effect of the factor treatment, but there was a crossover interaction. Density was significantly different in week one (1 nymphs/m^2^) against week two (0.2 nymphs/m^2^) (Estimate: −2.8875; Std. Error: 0.6285; P-value<0.05) and week three (Estimate: 1.2033; Std. Error: 0.3744; P-value<0.05). Nymph abundance decreased after tillage showing statistically differences among treatment and control. Treatments did not differ among them in the second week (Estimate: 1.3836; Std. Error: 1.1610; P-value=0.233), but they did in the third week (Estimate: −2.0871; Std. Error: 0.7814; P-value<0.05) being lower the density in treatment than in control. In plot B, initial density in control and treatment was the same (0.6 nymphs/m^2^) (Estimate: 0.3242; Std. Error: 0.4771; P-value= 0.497). After tillage, there were statistically differences among the treated cover plant rows in the second and third week (Estimate: −2.4563; Std. Error: 1.1022; P-value<0.05).

## 4. Discussion

Aim of controlling *X. fastidiosa* spreading is currently ongoing in the EU, considering that the management should be based on a combination of multiple tactics, such as vector monitoring, weed management, insecticides, natural enemies, repellents and pheromones [27, 51, 56]. Also, several protocols were developed to avoid short and long-range spreading of the pathogen [37]. In fact, one of the methods to decrease short-range spreading of *X. fastidiosa* is the control of the vectors such as nymphs or newly emerged adults (e.g., removal of ground vegetation) [29, 37, 51, 57].

In this study we assessed the effect of the management of the cover vegetation (i.e., mowing and tillage) in olive and vineyard organic orchards to decrease the density of nymphs of *X. fastidiosa* vectors. Our results indicated that tillage and mowing could be an efficient method for mechanical control since the nymphal density decreased between 10 to 50 % in the treatment plots compared to the control ones. The experiment was carried out according to nymphal seasonality described in López-Mercadal et al. [32], coinciding with the peak of nymphs between March and April. Also, nymphs were between N2 to N5 stage at the time of performing the treatment (26 % N2, 28 % N3, 18 % N4, 28 % N5). These stages are considered to be easiest to control due to their limited mobility compared to the highly mobile adults [29, 49–52].

We observed that nymphal density was reduced in 10-50 % when mowing and tilling weeds. These results are in line with Sanna et al., [29] that demonstrated that these techniques reduced *P. spumarius* density in a 20-60 %. Consequently, adult populations of vectors decrease and *X. fastidiosa* spreads lower. Also, Morris [53] observed that cutting cover decreased Auchenorrhyncha adult density in Arrhenatherum-dominated calcareous grasslands in the UK due to the management interfered hemipterans life cycle. Even so, cover management can contribute to eliminate the use of herbicides, fungicides, and pesticides [5, 54–55], that guarantees a sustainable production [5, 56], and prevents the immigration of new adults to shrubs and trees and decreases phytophagous activity in the crop, and then potential transmission of *X. fastidiosa* [59]. Hence, reducing the exposure of pesticides may contribute to the positive response of beneficial arthropod groups for the crop such as natural enemies [60]. For example, the spiders *Araniella cucurbitina* and *Synema globosum* that are dominant species in olive groves in Portugal have been reported as predators of *P. spumarius* [61]. Also, *Verralia aucta* (Pipunculidae) have been described as a parasitoid of the spittlebug in the United States [62].

In this study, management of the cover plants by mowing was usual in the olive plots, while tillage was the preferred technique in vine. No difference between the two techniques was detected in our study, but other authors showed that tillage can have a stronger effect than mowing on *P. spumarius* populations in Italy [29]. In addition, previous studies in Apulia showed that tillage performed in winter and spring reduced the abundance of *P. spumarius* and *N. campestris* on the cover vegetation and olive canopies [37]. In our case, we targeted nymphs only in the peak of the spring period [32] expecting to have a stronger effect on the population. Mowing or tillage during winter may have little effect due to the life cycle of the vectors, when most of the population is overwintering as egg. Farmers participating in our study performed mowing or tillage only once in each selected plot. It is expected that in agreement with Nickel and Hildebrandt [59], that a more severe regime of mowing would have higher impact causing a long-term exclusion of Auchenorrhyncha species. However, one limitation of the management of plant cover in crops is that plants remaining in the border of orchards and near the tree may also hold an important number of nymphs and its management would be also required [29]. Besides, these practices have been proven to also decrease other taxons such as spiders [50, 63–64], orthopterans [65–66], lepidopterans [67] and coleopterans [63] because grasslands act as refuges for these auxiliar fauna.

It is important to point out that permanent removal of ground vegetation is very common in traditional agricultural management in the Mediterranean Basin. Plant cover removal is usually performed to avoid competence for water resources and to facilitate crop management (i.e, use of machinery, fruit picking, etc.). However, it is known that total removal of plant cover would imply to serious ecological problems and enhance pests in the crop, since possible reservoirs of natural enemies are eliminated [68–70]. In addition, total removal of plant cover may also lead to serious soil loss as documented in several regions of Spain [71].

It is relevant to note that *P. spumarius* and *N. campestris* do not act as pests in crops, but their population is conditioned by the management intensity of each plot [58] because they have two ground-dependent stages: nymphs and egg laying. As commented before, control of eggs is not feasible because they are laid in the stubble during autumn. Therefore, specific tillage or mowing of cover plants is advised in specific period of time to eliminate development and feeding sites for vectors. For this, and taking into account its life cycle, early stage nymphs are the most vulnerable to biotic and abiotic factors due to its low mobility and thinner spittle [72]. Consequently, mechanical control methods are limited to weeks where nymphs are in N1 and N2 stages. Meanwhile nymphs from N3 to N5 gain mobility and can reach other plants then reducing the efficacy of mowing or tillage. In this study, we did not assess the adult population after conducting the mechanical control of nymphs and this needs further exploration. In Italy, it was reported that reducing nymphal density would decrease adult vector densities [29].

In conclusion, we showed that spring mowing and tilling of the plant cover vegetation in Mediterranean olive and vineyard organic farming crops significantly reduced the nymphal density of vectors of *X. fastidiosa*. It is needed to further explore the impact of mechanical control of nymphs on vector adult population, as well as the effect on *X. fastidiosa* risk of transmission.

## Author Contributions

Conceptualization, MAM and JLM; Methodology, MAM and JLM.; Formal Analysis, JLM; Investigation, MAM, JLM and PM; Writing – Original Draft Preparation, JLM; Writing – Review & Editing, MAM, JLM and PM; Funding Acquisition, MAM.

## Data Availability Statement

“Not applicable”

## Acknowledgments

We thank the farmers for their participation and let as conduct the experiment.

